# The gut-liver axis and the role of bacteria in severe dengue virus infection in mice

**DOI:** 10.64898/2026.09.21.749971

**Authors:** Adriana Pliego-Zamora, Jaehyeon Kim, Naphak Modhiran, Karli Takizawa, Ennae Gollasch-Miller, Ran Wang, Paul R. Young, Daniel Watterson, Sumaira Z. Hasnain, Parimala R. Vajjhala, Yuji Sekiguchi, Helle Bielefeldt-Ohmann, Dieter M. Tourlousse, Katryn J. Stacey

## Abstract

Patients with severe dengue virus (DENV) infection can experience hemorrhage, shock and organ damage that coincides with defervescence and declining viremia. In addition to vascular leak, published work has shown that gut barrier permeability and elevated serum lipopolysaccharide (LPS) levels correlate with DENV disease severity. We previously described profound gut pathology in a mouse model of infection. This led to the hypothesis that a loss of gut epithelial barrier integrity and the influx of microbial products contribute to exacerbated inflammation and severe dengue disease. Consistent with this, here we demonstrated that DENV2 infection of AG129 mice compromised the gut epithelial barrier, promoted bacterial translocation, and induced microbial changes indicative of dysbiosis, including a reduced abundance of putatively beneficial microbial taxa and disrupted microbial circadian oscillations. Depleting gut bacteria with antibiotics from 1-day post-infection sharply reduced gut pathology as well as viral RNA in the proximal colon and liver, and the number of infected Kupffer cells in the liver. There was no evidence for systemic suppression of viral replication. We propose that viral infection of intestinal macrophages leads to inflammatory damage and mucosal barrier breakdown, dysbiosis and gut leak, with gut bacteria-dependent products promoting the susceptibility of Kupffer cells to DENV infection. Given the importance of liver damage in severe and fatal dengue disease, therapies that protect the gut barrier may help reduce disease severity.

## Introduction

Dengue virus (DENV) is an enveloped, sense-strand RNA virus of the *Flaviviridae* family, transmitted to humans by mosquitoes and is responsible for approximately 100 million symptomatic infections annually [1]. There are four DENV serotypes, DENV1 to DENV4 with antibody-dependent enhancement of infection proposed to contribute to exacerbation of disease upon subsequent infection with a different serotype. Clinical manifestations range from asymptomatic infection to uncomplicated dengue fever, to the life-threatening dengue hemorrhagic fever (DHF) and dengue shock syndrome (DSS). The severe pathology of DENV infection involves plasma leak leading to shock, fluid accumulation and respiratory distress, or hemorrhage associated with thrombocytopenia, or severe organ involvement including signs of liver damage [2, 3]. In 2009 the WHO introduced categorization of DENV infection as dengue with or without warning signs, and severe dengue. The warning signs include several gastrointestinal and hepatic manifestations including abdominal pain, persistent vomiting, gastrointestinal bleeding, and liver enlargement [3].

Elevated serum lipopolysaccharide (LPS) levels correlate with disease severity in patients [4, 5] suggesting that epithelial barriers may be compromised in DENV infection. The intestinal epithelium is a critical barrier separating the luminal microbiota from the systemic circulation. Disruption of this barrier, or “gut leak”, permits translocation of bacteria and microbial products, including LPS, which can amplify systemic inflammation, endothelial activation and organ injury in a range of diseases [4–7]. Although several studies have reported evidence of intestinal barrier dysfunction during severe dengue [5, 6], it remains unclear whether gut leak actively contributes to disease progression or simply reflects advanced systemic inflammation.

Importantly, gut leak has been demonstrated in patients using the lactulose to mannitol excretion ratio method (LMER), and determined to correlate with dengue severity [5]. For patients who developed severe disease, 24/27 had a positive LMER test, compared with only 7/59 non-severe patients. In that study, the levels of circulating LPS and fungal (1→3)-β-D-glucan positively correlated with the LMER results, in the absence of live bacterial isolation from the blood of the total 86 cases [5, 6]. Evidence of gut damage in DENV-infected patients has also been provided by elevated circulating intestinal fatty acid binding protein (I-FABP), which is normally found in intestinal epithelial cells and is released with tissue damage [6]. Moreover, studies of bacterial coinfection in dengue have noted the presence of gut commensal organisms [8]. Bacteremia was observed in 0.18-1.2% of DENV infections overall, 7% of DHF/DSS cases and in 14-44% of fatal cases [9]. In mouse models, DENV infection is observed in intestinal macrophages [10–12] accompanied by extensive mucosal inflammatory pathology including depletion of goblet cell mucus content [11–13].

We hypothesize that severe DENV infection can cause gut barrier impairment promoting the translocation of bacteria and/or their products to the liver, further exacerbating the pathology of dengue disease. Therefore, we examined DENV-infected mice for signs of gut barrier breakdown and bacterial entry and assessed the role of bacteria in the development of gut pathology and viral replication in AG129 mice using antibiotic treatment. The results reveal DENV-induced gut leak and dysbiosis and, unexpectedly, suggest a role for gut bacteria in the susceptibility of Kupffer cells to infection, late in the disease course.

## Methods

### Cell culture

Vero cells were cultured as previously reported [11] in Dulbecco’s modified Eagle’s medium (DMEM) supplemented with 10% heat-inactivated fetal bovine serum (FBS), 2 mM L-alanyl-L-glutamine dipeptide, 50 U/mL penicillin, and 50 µg/mL streptomycin (‘complete DMEM’). Mosquito C6/36 cells were cultured in RPMI medium supplemented with 10% FBS, 2 mM L-alanyl-L-glutamine dipeptide, 50 U/mL penicillin, 50 µg/mL streptomycin, and 25 mM HEPES, and maintained in sealed flasks at 28°C in an incubator without humidity or CO_2_ control. All reagents were from Gibco.

### Dengue virus

DENV2 mouse-adapted strain D220 [14] was kindly provided by Prof. Eva Harris (School of Public Health at the University of California, Berkley, USA). Viral stocks were obtained following a previously reported protocol [11]. Briefly, virus was grown in C6/36 cells, and viral particles were concentrated by ultracentrifugation with a 20% sucrose cushion using a SW 32 Ti rotor (Beckman Instruments). Viral pellets were resuspended in phosphate buffered saline (PBS) overnight at 4°C and aliquots were stored at -80°C.

### Virus titration

Fluorescent immuno-plaque assay was performed to quantify infectious viral particles in the DENV D220 stock and serum samples [11, 15]. Briefly, Vero cells were seeded in 96 well plates and incubated with 10-fold serial dilutions of the samples in serum-free Opti-MEM. The inoculum was then removed and M199 medium (Thermo Fisher) containing 2% FBS, 50 U/mL penicillin, 50 µg/mL streptomycin, 1.5% carboxymethylcellulose and 25 mM sodium bicarbonate was overlaid on the cells. Plates were incubated for 4.5 days and then immunolabeled using human anti-DENV E antibody (4E11) [16] and goat anti-human IgG NIR800 (Li-Cor Biosciences). Immunolabeled plates were scanned on the Odyssey Imager (Li-Cor Biosciences) in the 800 nm channel. The titer was calculated by counting fluorescent plaques and determined as focus-forming units per mL (FFU/mL).

### NS1 ELISA

Concentrations of NS1 in serum samples were determined by a sandwich ELISA as previously reported [11, 17]. Microlon high binding 96 well plates (Grenier) were coated with capture anti-NS1 Gus2 antibody [18] and then blocked with 1% BSA in PBS. A standard curve was included in each 96 well plate using purified insect cell-derived NS1 [19]. Standards and 2-fold dilutions of serum samples were incubated on the plate, then washed with PBS-T before protein G-purified polyclonal rabbit anti-NS1 serotype 2 antibody was added and incubated. The plate was washed, incubated with HRP-conjugated goat anti-rabbit antibody (Cell Signaling Technology), washed and the reaction was developed with 3,3’,5,5’-tetramethylbenzidine (Sigma) and H_2_SO_4_ was added as stop solution. Optical density at 450 nm was measured using a SpectraMax 190 (Molecular Devices).

### Animals and tissues

Research was conducted with University of Queensland (UQ) animal ethics committee approvals (016/21 and 185/24). Mice were bred in-house and housed at a UQ Biological Resources animal facility under specific pathogen-free conditions, with 12:12 h day-night cycle. Male and female AG129 mice, which lack receptors for type I and II interferon (IFN-α/β/γ) on a 129-strain background, were infected intraperitoneally (*i.p*) at 5-10 weeks of age with 1×10^5^, 3×10^5^ or 5×10^5^ FFU of DENV D220 in 100 µL of PBS. Mock infected mice received 100 µL of PBS *i.p*. Mice were monitored daily for up to five days. Clinical scores were determined by observers blinded to the groupings, who assessed signs of disease including locomotion, appearance, and behavior. Fresh feces were assessed daily and diarrhea was scored using a previously published protocol [11] described as 0: solid and bouncy, 1: solid but easily crushed, contains undigested food contents, 2: semi-liquid state, containing < 50% liquid and 3: viscous yellow liquid. Mock and infected mice were euthanized with CO_2_ at 4 days post infection (d.p.i) or whenever they reached humane end-points as per ethics approval. Blood was obtained via tail bleed at 3 d.p.i or by heart puncture at time of euthanasia. Samples of stomach, small intestine, large intestine, liver, spleen, kidney, lung and bone marrow were collected in RNAlater (Thermo Fisher) and stored at -80°C. In some experiments mice were perfused with PBS before tissue collection. Tissues for histology were fixed in 10% neutral buffered formalin for 3 days and then transferred to 80% ethanol.

### RNA extraction and qRT-PCR

RNA extraction and qRT-PCR were done as previously described [11]. Briefly, tissues were stored in RNAlater (Thermo Fisher), and RNA was extracted using TRIzol Reagent (Life Technologies), stainless-steel beads (Qiagen) and TissueLyser II (Qiagen). RNA concentrations were determined using a NanoDrop OneC Microvolume UV-Vis Spectrophotometer (Thermo Scientific). Four µg of RNA was reverse transcribed, using random hexamers (IDT) and Superscript IV (Invitrogen). cDNA samples were diluted 1:10 with Ultra Pure water (Invitrogen) and 2 µL of the diluted samples were used for qRT-PCR in 384 well plates using QuantiNova SYBR^TM^ Green PCR Master Mix (Thermo Fisher) with primers (Table 1). Samples were run in triplicate in a QuantStudioTM 6 Real Time PCR system (Life Technologies). Expression was determined relative to the *Hprt* gene [relative expression = 2^(Cт(Target gene) – Cт(Hprt))^] (38). The average Cт values for the *Hprt* gene were not affected by infection.

### Depletion of bacteria with antibiotics cocktail

Antibiotics solutions were prepared 24 hours before each experiment and contained between one and four antibiotics [20]. Ampicillin (1 mg/mL), vancomycin (1 mg/mL), metronidazole (0.5 mg/mL), and neomycin (1 mg/mL), were dissolved in water supplemented with 10% sucrose, and filtered through a 0.22 μm filter. Antibiotic solutions were aliquoted into mouse cage water bottles on day 1 post-infection (1 d.p.i.). Untreated mice received water supplemented with 10% sucrose. Daily intake of antibiotics was assessed by weighing the water bottles.

### Fecal sample collection and DNA extraction

The microbiome across mice was synchronized for at least two weeks before infection, as described above. The bedding of every cage was collected, mixed, and re-distributed to all cages every second or third day. Every 12 hours from the evening prior to infection fecal pellets were collected directly into 2 mL screw cap tubes, and stored at -80°C. Total DNA was extracted from whole fecal pellets weighing between 50-200 mg. Pellets were first homogenized using the DNeasy 96 Powersoil Pro QIAcube HT Kit (Qiagen) following the manufacturer’s protocol. Briefly, CD1 buffer and 1 mm diameter glass beads (BioSpec Products) were used to homogenize the sample by bead beating using the Powerlyser 24 homogenizer (Mo-Bio). Samples were heated and shaken at 65°C for 10 minutes at 1000 RPM, on the Eppendorf ThermoMixer (Eppendorf) and then centrifuged for one minute at 10 000*g*. The resulting lysates were transferred to a 96 deep well plate (Axygen), then frozen and thawed to further lyse the cells. CD2 buffer was added to the thawed lysate and then centrifuged at 10 000*g*. The resulting supernatant was transferred to a new S block (supplied in the kit) and DNA extraction was performed using the Qiagen QIAcube HT instrument (Qiagen) according to the manufacturer’s protocol. Final elution volume was 80 µL.

### Quantification of fecal bacteria

Fresh fecal samples were collected directly from the anus into sterile pre-weighed tubes at the designated time points. Sterile PBS was added at 1 µL/mg of feces, fecal pellets disrupted by vigorous pipetting and then left at RT for 10 min. Fecal solutions were serially diluted 10-fold in PBS and 10 µL of each dilution (10^-1^ to 10^-8^) were spotted on horse blood agar plates (Edwards). For each experiment, plates were incubated in the indicated chambers in the figure legend at 37°C for 3 days. Data is expressed as colony formation units (CFU) per mg feces.

### Full-length 16S rRNA gene amplicon sequencing

Near full-length 16S rRNA gene amplicon libraries were generated using a two-step tailed PCR strategy. First-round PCRs contained 1× KAPA HiFi HotStart ReadyMix, 500 nM each of forward primer (5’-TTTCTGTTGGTGCTGATATTGC<u>AGRGTTYGATYHTGGCTCAG</u>-3’; the underlined region represents the bacterial ‘27F’ 16S rRNA gene primer) and reverse primer (5’-ACTTGCCTGTCGCTCTATCTTCTA<u>CGGYTACCTTGTTACGACTT</u>-3’; the underlined region represents the bacterial ‘1492R’ 16S rRNA gene primer), and 10 ng of template DNA. Thermal cycling conditions were: 98 °C for 3 min; 18-22 cycles of 98 °C for 10 s, 55 °C for 10 s, and 72 °C for 90 s; and 72 °C for 5 min. Amplicons were purified using the Agencourt AMPure XP system with a 0.6× bead-to-sample ratio. Barcode sequences were then added using the PCR Barcoding Expansion 1–96 kit (EXP-PBC096) in a second round of PCRs, containing 1× KAPA HiFi HotStart ReadyMix, 250 nM PCR barcode (BC01–96), and approximately 1 ng of purified first-round PCR product. Thermal cycling conditions were: 98 °C for 3 min; 8 cycles of 98 °C for 10 s, 60 °C for 10 s, and 72 °C for 90 s; followed by a final extension at 72 °C for 5 min. Barcoded amplicons were purified using AMPure XP beads as above, quantified with a Qubit dsDNA High Sensitivity Assay kit, and pooled at equimolar concentration. Subsequent DNA end prep and adapter ligation were performed using the Ligation Sequencing Kit 14 (SQK-LSK114) according to the manufacturer’s instructions. Sequencing was conducted on a MinION Mk1D device using an R10.4.1 flow cell. Basecalling was performed with Dorado v1.1.1 (available from https://github.com/nanoporetech/dorado) using the super-accuracy model (dna_r10.4.1_e8.2_400bps_sup@v5.2.0). After conversion to FASTQ format and removal of sequence reads longer than 2,250 bp, demultiplexing was performed using Dorado’s demux command with options --emit-fastq --no-trim --kit-name EXP-PBC096. Sequence reads then underwent primer trimming using Cutadapt v4.9 (Martin et al., 2019), with the options -g ’AGRGTTYGATYHTGGCTCAG…AAGTCGTAACAAGGTARCCG’ -e 3 -O 20 -m 1250 -M1750 --max-n 0 --rc --discard-untrimmed, followed by removal of sequence reads with a mean quality score below 16 using fastq-filter v0.3.0 (available from https://github.com/LUMC/fastq-filter). To minimize the impact of uneven sequencing depth, processed sequence reads were randomly subsampled to 40,000 reads per sample using seqtk v1.5 (available from https://github.com/lh3/seqtk) prior to species-level taxonomic profiling with Emu (see below).

### Microbiome community profiling and analysis

Microbial community profiling was performed using Emu (v3.5.4; [21]) against the “SBDI Sativa curated 16S GTDB database”, built from the Genome Taxonomy Database (GTDB; R10-RS226 [22]. The strictly filtered version of the database (“sbdi-gtdb-sativa.r10rs226-2”, with 5 genomes per species) was obtained from [23] and formatted using Emu’s build-database command. Species-level abundances were estimated with Emu’s abundance command and subsequently aggregated at the genus level if required, retaining estimated read counts for downstream analyses.

All subsequent analysis was performed in R (v4.5.1; [24]). Bray-Curtis dissimilarities were calculated using vegan’s (v2.7; [25]) vegdist function (method=”bray”, binary=FALSE), and principal coordinate analysis (PCoA) performed using ape’s (v5.8;[26]) pcoa function with correction=”lingoes”. Differential abundance analysis was performed using linear models for compositional microbiome data, as implemented in the R package LinDA (v0.2.0; [27]), with options adaptive=FALSE, imputation=FALSE, pseudo.cnt=0.5. Here, we restricted analysis to species detected in >20% of samples, and tests were considered significant at an adjusted P value of 0.05. Two types of mixed-effect linear models were fitted to: (i) compare species abundances between pre- and post-infection samples, for each post-infection timepoint and experimental group separately [model formula: counts∼timepoint+(1∣ animal)], and (ii) compare species abundance trajectories between experimental groups [model formula: counts∼timepoint_numeric∗experimental_group+(1∣ animal)].

### Vascular leakage

Evans Blue dye at 5 mg/mL in PBS (Sigma Aldrich) was sterilized with a 0.22 µm filter, and 200 µL injected *i.v.* into mock and infected mice and allowed to circulate for 2 hours before mice were euthanized and perfused. Briefly, mice were anaesthetized with isoflurane, then the heart was exposed, a cut was made in the right auricle and 40 mL of PBS was pumped through the left ventricle. Tissues of interest were collected in pre-weighed tubes, 1 mL of N, N-dimethylformamide was added and samples incubated at 37 °C for 24 hours. Extracted dye in the supernatant was assessed by measuring OD_620_ and quantified using a standard curve. Data is expressed as ng of Evans Blue dye per mg of tissue weight.

### Gut leak

Mock and infected mice received 300 mg/kg of 4 kD FITC-Dextran (Sigma) in water by oral gavage. Oral gavage control mice received 120 µL of water. After 4 hours post-gavage, mice were euthanized with CO_2_, and blood was collected by heart puncture. Serum was recovered and used to quantify 4 kD FITC-Dextran by measuring fluorescence in a Clariostar plate reader (BMG BIOTECH) using a standard curve of 4 kD FITC-Dextran. Data is expressed as µg/mL of serum.

### Bacteria culture from tissues

After mice were euthanized, liver and mesenteric lymph nodes (MLN) were harvested under sterile conditions and placed into pre-weighed tubes containing 200 µL of sterile 0.5% Igepal (Sigma) in PBS. Tissues were homogenized using a TissueLyser II (Qiagen) and sterile stainless-steel beads (Qiagen). Homogenized tissues were diluted 1:10 with sterile PBS and 30 µL of each tissue dilution was spread onto two 5% horse blood (Thermo Fisher) agar plates. Plates were placed in anaerobic (Thermo Fisher) or microaerophilic (Biomerieux) chambers at 37°C for 4 days. Data is expressed as colony formation units (CFU) per 100 mg of liver or per whole MLN.

### Histology and pathology scoring

Formalin-fixed tissues were routinely processed for paraffin-embedding at the UQ School of Veterinary Science (SVS) histology laboratory. Sections (4 µm) of small intestine Swiss rolls and large intestines were stained with hematoxylin and eosin (H&E) and examined microscopically by a veterinary pathologist blinded to the treatments as previously described [11]. Detection of the DENV NS3 was performed by immunohistochemistry as previously reported [11]. Warthin-Starry silver stain was performed as previously reported [28].

### Flow cytometry

Liver leukocytes were isolated from perfused tissues by centrifugation [29]. Livers were finely chopped in 10 mL digestion solution containing 1 mg/mL Collagenase IV (Gibco), 0.4 mg/mL Dispase II (Sigma Aldrich) and 20 μg/mL DNAse1 (Roche) in 1X HBSS (Gibco) and incubated at 37 °C for 45 minutes on a rocking platform before mashing through a 70 μm filter (Falcon). Hepatocytes were extracted by one run of a slow spin (50*g*) and the non-parenchymal cells (NPCs) in the supernatant were then pelleted (500*g*), resuspended in Percoll (Cytiva) and centrifuged at 700*g* with no brake. The supernatant containing hepatocytes was discarded and the enriched leukocyte pellet was resuspended in ACK lysis buffer (Gibco). Cells were washed once in PBS and once in staining buffer (SB, PBS with 2% FBS), and the pellet was resuspended in SB. One million cells were immunolabeled in FACS tubes, first for 15 min at RT with Zombie violet dye (Biolegend), and then for 30 min on ice with antibodies against cell surfaces markers CD45-AF700, CD11b-FITC, F4/80-PE594, Ly6C-PE and Tim4-PECy7 (all from Biolegend) in SB containing 2.4G2 hybridoma supernatant (10%) to block Fc receptors. Cells were washed with cold SB, resuspended in permeabilization buffer (PBS with 0.1% saponin) followed by intracellularly immunolabeling with primary antibody goat anti-NS3 (in-house) for 30 min on ice, washed with cold SB, and then secondary donkey anti-goat AF647 (Abcam) in permeabilization buffer was added for 20 min on ice. Cells were washed and resuspended in cold SB, counting beads were added and samples analyzed using a Cytoflex flow cytometer (Beckman Coulter LifeSciences). Single-color controls were used for compensation, and unstained and fluorescent minus one samples were used to confirm gating strategies.

### Data availability

Raw sequencing data have been deposited in NCBI’s Sequence Read Archive under BioProject PRJNA1405243.

### Software

Statistical analyses were performed using GraphPad Prism 9 software (GraphPad Software Inc, San Diego, CA, USA). Error bars represent the SD. Flow cytometry data was analyzed with Flow Jo (Tree Star). The microbiome data was analyzed using R as described above. Some diagrams were created in Biorender https://bioRender.com.

## Results

### DENV infection can cause gut permeability and bacterial translocation

Following from our prior findings of profound gut inflammatory pathology and loss of mucus barrier in mouse models of severe DENV infection [11], we further investigated if DENV infection can cause gut permeability, similar to that documented in severe dengue patients [5]. AG129 mice were infected with DENV and orally administered 4 kDa FITC-dextran, which normally has limited intestinal translocation, at various times post infection. We observed a sudden increase of serum FITC-dextran between 3.2 and 4.2 d.p.i. (76 and 100 hours post-infection [h.p.i.]) with onset during peak viremia, and while serum NS1 concentrations were still increasing (Fig. 1A). Plating liver and mesenteric lymph node (MLN) homogenates and growth under anaerobic or microaerophilic conditions revealed bacterial translocation from 4.2 d.p.i (100 h.p.i) (Fig. 1A). Replicate experiments confirmed late gut leak (Supp. Fig. 1A-C) and bacterial translocation into the liver (Supp. Fig. 1A and B). Whilst gut permeability and bacterial influx were detected in several experiments, they were not always observed. In some experiments, bacterial translocation into the liver was not detected and this correlated with either lack of gut leak or its later onset (Supp. Fig. 1C and D). A further experiment showed gut leak at 4.2 d.p.i occurring in a virus dose-dependent manner (Fig. 1B). In line with the previously observed depletion of goblet cell mucus [11], we found that mRNA expression of the major mucin gene *Muc2* was reduced in the proximal colon, as well as the goblet cell transcription factors *Atoh1* and *Spdef*, with a trend for virus dose-dependency (Fig. 1C). To further investigate bacterial translocation we examined colon sections using a Warthin-Starry silver stain. This showed the absence of bacteria in the proximal colon crypts of uninfected mock mice, as expected (Fig. 1D). However, in infected mice bacteria were seen in the crypts, traversing the epithelial layer at the base of the crypt, and penetrating further into the submucosa.

**Figure 1.**
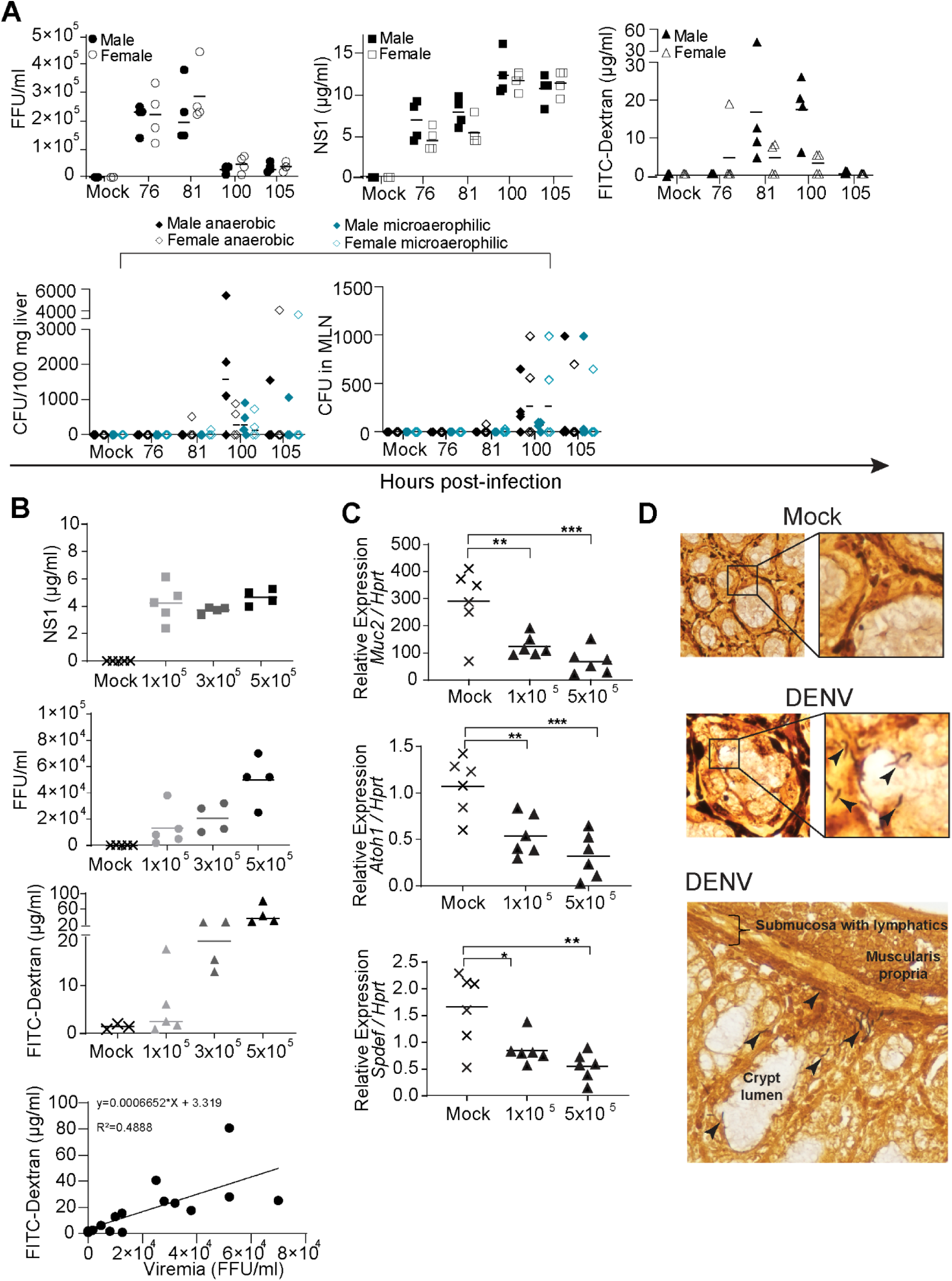
Mouse DENV infection compromises gut integrity and allows bacterial tissue penetration. Groups of 6–7-week-old male and female AG129 mice were infected *i.p.* with 3×10^5^ FFU DENV2. **A.** Viremia, levels of NS1, gut leak measured by detection of serum FITC-dextran following oral gavage, and bacterial translocation to the liver and MLN were assessed at the indicated time points, n = 4 per group. Bacterial counts were assessed under anaerobic and microaerophilic culture conditions, expressed per 100 mg liver or for total MLN homogenate. **B.** Male mice were infected with 1×10^5^, 3×10^5^ or 5×10^5^ FFU DENV2. Levels of NS1, viremia and gut leak were assessed at 4 d.p.i (96 – 100 h.p.i), n = 4-5 per group. Linear regression is shown for viremia correlation with gut leak. **C.** Expression of mRNAs for *Muc2* and goblet cell transcription factors *Atoh1* and *Spedf* from proximal colon samples at 4 d.p.i from mock or infected AG129 mice with the indicated doses of DENV2, n = 6 per group from two separate experiments. **D.** AG129 mice were infected with 1×10^5^ FFU DENV2 and GI tissues were harvested at 4 d.p.i.. Warthin-Starry silver stains of colon. Black arrowheads indicate bacteria seen in tissues and crypt lumens in 4/5 infected mice but not in mock-infected mice. N.B. Some nuclei stain black. Lines represent the mean, except for the CFU where it represents the median. \**p*<0.03, \*\**p*<0.002 and \*\*\**p*<0.0002 using one-way ANOVA. CFU, colony forming units.

### DENV infection causes dysbiosis

Gut inflammation and leak are frequently associated with dysbiosis [30, 31]. We assessed the microbiome of DENV-infected mice using fecal samples over a time course before and after infection using Nanopore long-read sequencing of near full length 16S rRNA gene amplicons. Mice were infected at 8 am, and fecal samples taken every 12 hours on most days with blood sampling at 3 d.p.i. (Fig. 2A). Evening samples (−0.5, 1.5, 2.5 and 3.5 d.p.i) were used to determine the composition of the microbiome as late post-infection as possible before diarrhea started at 4 d.p.i.. Based on unconstrained ordination analysis (PCoA of species-level Bray– Curtis dissimilarities), we observed that DENV infection induced greater changes in the gut microbiome than mock treatment, as reflected by the larger shift along the first PCoA axis (Fig. 2B). Consistent with this, Bray–Curtis dissimilarity from the pre-infection microbiome increased more strongly in DENV-infected mice, becoming significant from 2.5 d.p.i. onwards (Fig. 2C). The increase observed in mock controls between 2.5 and 3.5 d.p.i. may reflect stress associated with blood sampling at 3 d.p.i., which could have altered the gut microbiome. Genus-level differential abundance analysis, performed separately for each post-infection time point and treatment group, likewise identified substantially more genera with altered abundance relative to pre-infection levels in DENV-infected mice than in mock controls at both 2.5 and 3.5 d.p.i. (Fig. 2D). In addition, a linear model incorporating treatment, time, and their interaction identified several genera whose temporal abundance trajectories differed significantly between DENV-infected and mock-treated mice (Fig. 2D). Genera showing a greater decline following DENV infection included *Lactobacillus*, *Limosilactobacillus*, and UBA7173, whereas *Scatocola*, *Anaerotignum*, UBA11957, MGBC122484, and *Acetatifactor* exhibited greater increases (Fig. 2D and Supp. Fig. 2). Finally, comparisons of evening and morning samples indicated that DENV infection disrupted circadian rhythms in the gut microbiome (Supp. Fig. 3). For multiple genera, including *Paramuribaculum, Petralouisia* and CAG-873, morning abundances in DENV-infected mice shifted towards pre-infection levels observed in evening (fasted) samples. While such changes may partly reflect the reduced food and water intake associated with infection, this did not explain all microbiome changes as a range of genera including UBA11957, COE1 and *Acutalibacter* had the opposite effect, with evening abundances moving towards the pre-infection morning levels.

**Figure 2.**
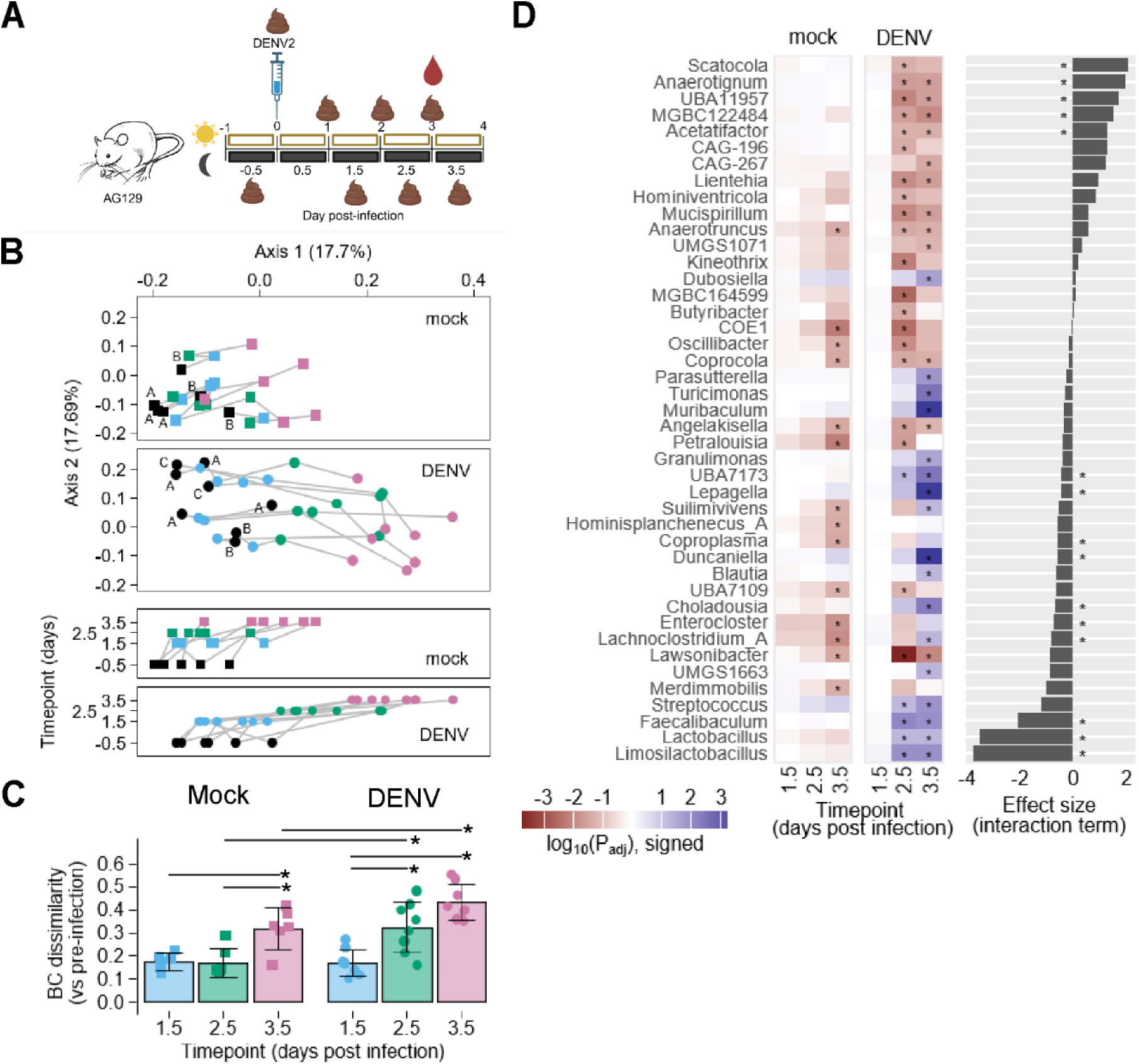
Mouse DENV infection causes dysbiosis and interruption of gut microbial diurnal rhythm. AG129 mice were infected *i.p.* with 1×10^5^ FFU DENV2. **A.** Fecal samples were collected before and after DENV2 infection, at 8 am and 8 pm. (Figure was created in BioRender https://bioRender.com). **B.** Principal coordinates analysis (PCoA) of fecal microbiome samples based on species-level Bray-Curtis (BC) dissimilarities. Samples from mock-treated (n = 6) and DENV-infected mice (n = 8) are shown in separate panels for clarity, although the ordination was computed using the full dissimilarity matrix. Each symbol represents an individual sample and is colored by collection timepoint. Grey lines connect longitudinal samples from the same mouse, and letters adjacent to the pre-infection samples (in black) indicate cage assignments. The two lower panels show sample coordinates along the first principal coordinate at different times. **C.** Community-wide changes in microbiome composition, quantified as species-level BC dissimilarities relative to animal-matched pre-infection samples. Bars indicate mean values, error bars represent the standard deviation, and colored symbols denote individual data points. Pairwise comparisons were performed using the Mann–Whitney U test and considered statistically significant at an unadjusted two-sided P value < 0.05. **D.** Heatmap summarizing differential abundance results of specific genera at each timepoint, shown separately for each experimental group. Fill colors represent signed adjusted P values, and asterisks indicate significant changes (adjusted P value < 0.05 and |log fold change| > 1). The adjacent bar plot shows the effect size of the treatment–time interaction term from a linear model including timepoint (as a continuous variable) and treatment group; significant effects (adjusted P value < 0.05) are marked by an asterisk.

**Figure 3.**
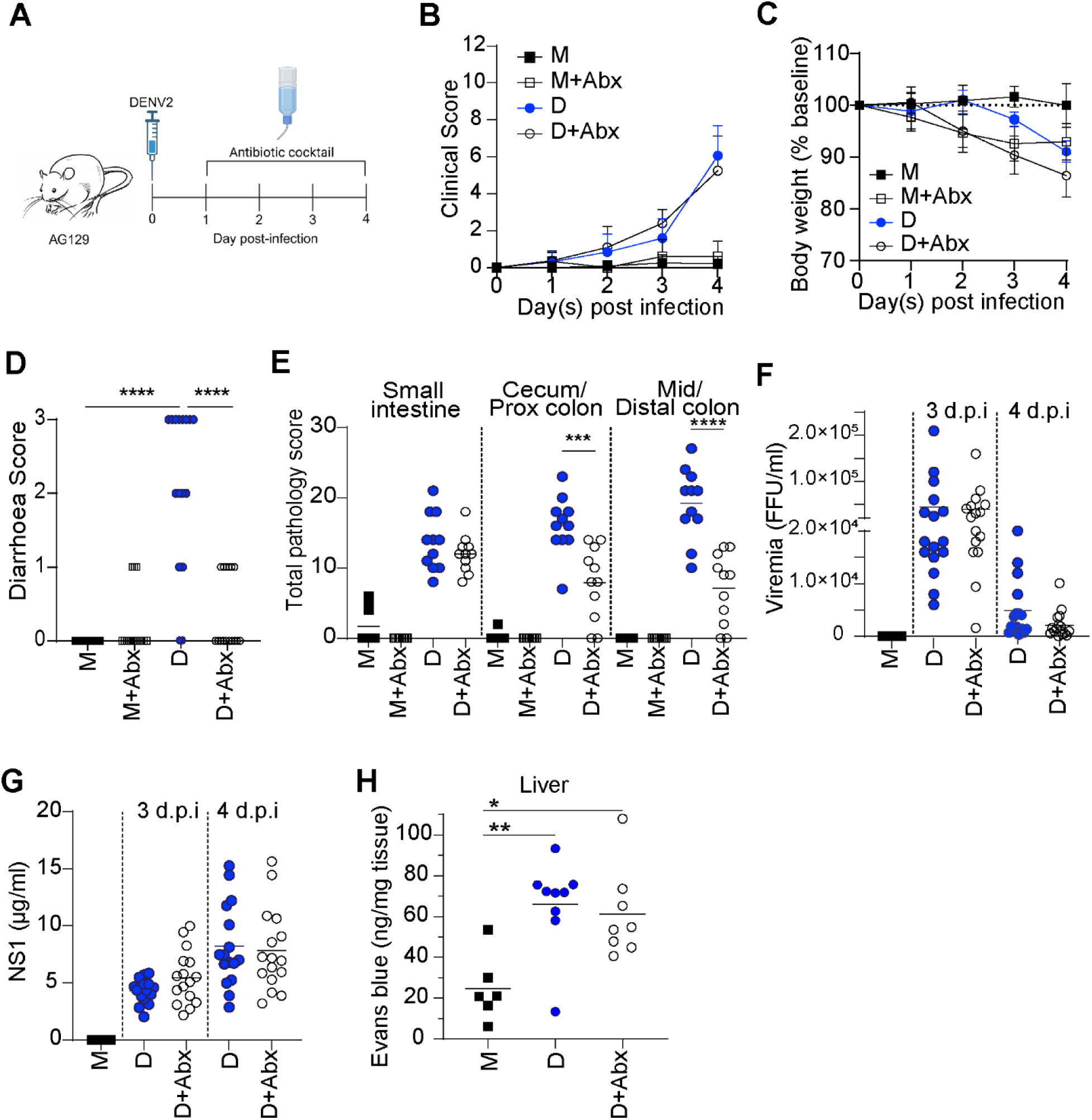
Depletion of bacteria with a broad spectrum antibiotic cocktail treatment reduces gastrointestinal tract DENV-associated pathologies without reducing viremia. **A.** Eight to 10-week-old male AG129 mice were *i.p.* infected with 1×10^5^ FFU DENV2. A cocktail of antibiotics (ampicillin, vancomycin, metronidazole and neomycin) was given in the drinking water from 1 d.p.i. until the end of the experiment (Figure was created in BioRender https://bioRender.com). B. Clinical scores, C. weight loss, D. diarrhoea incidence at 4 d.p.i, E. gut pathology scores at 4 d.p.i., F. viremia, and G. levels of NS1 in serum. For panels B-G, n = 11-16 per group from two experiments for M+Abx, and from three experiments for all other groups. H. Vascular leak in the liver at 4 d.p.i. assessed by tissue content of Evans blue after i.v. administration, n = 6-9 per group from two experiments. Lines represent the mean, \**p*<0.05, \*\**p*<0.01, \*\*\**p*<0.001, \*\*\*\**p*<0.0001 of unpaired *t*-test. B and C are the mean ± SD. M, mock-infected; D, DENV-infected; Abx, antibiotics cocktail-treated, FFU, focus-forming unit.

### Bacterial depletion reduces colon inflammation during DENV infection

We examined the effect of bacterial depletion on disease presentation. Male AG129 mice were infected with DENV and treated with a cocktail of 4 antibiotics in the drinking water from day 1 post-infection (Fig. 3A). The combination of ampicillin, neomycin, metronidazole and vancomycin has been shown to profoundly reduce bacteria in feces, with approximately 3-log reduction in 16S rRNA copies per fecal pellet [20, 32, 33]. Antibiotic treatment reduced numbers of fecal bacteria that could be cultured on blood agar under microaerophilic conditions by at least 3-4 log units after 1 day of treatment (Supp. Fig. 4A). The aggressive antibiotic treatment itself compromised mouse wellbeing to some extent, as uninfected antibiotic-treated mice showed signs of loss of condition and suffered a significant weight loss over a 3-day period (Fig. 3B-C). Published work suggests that this could be expected, as loss of microbiota with antibiotics causes metabolic changes that result in loss of fat, liver and muscle tissue [32–34]. With DENV infection, there was no difference in clinical scores between antibiotic-treated and untreated mice, although antibiotic-treated mice lost more weight, consistent with the effect of antibiotics alone (Fig. 3B-C).

**Figure 4.**
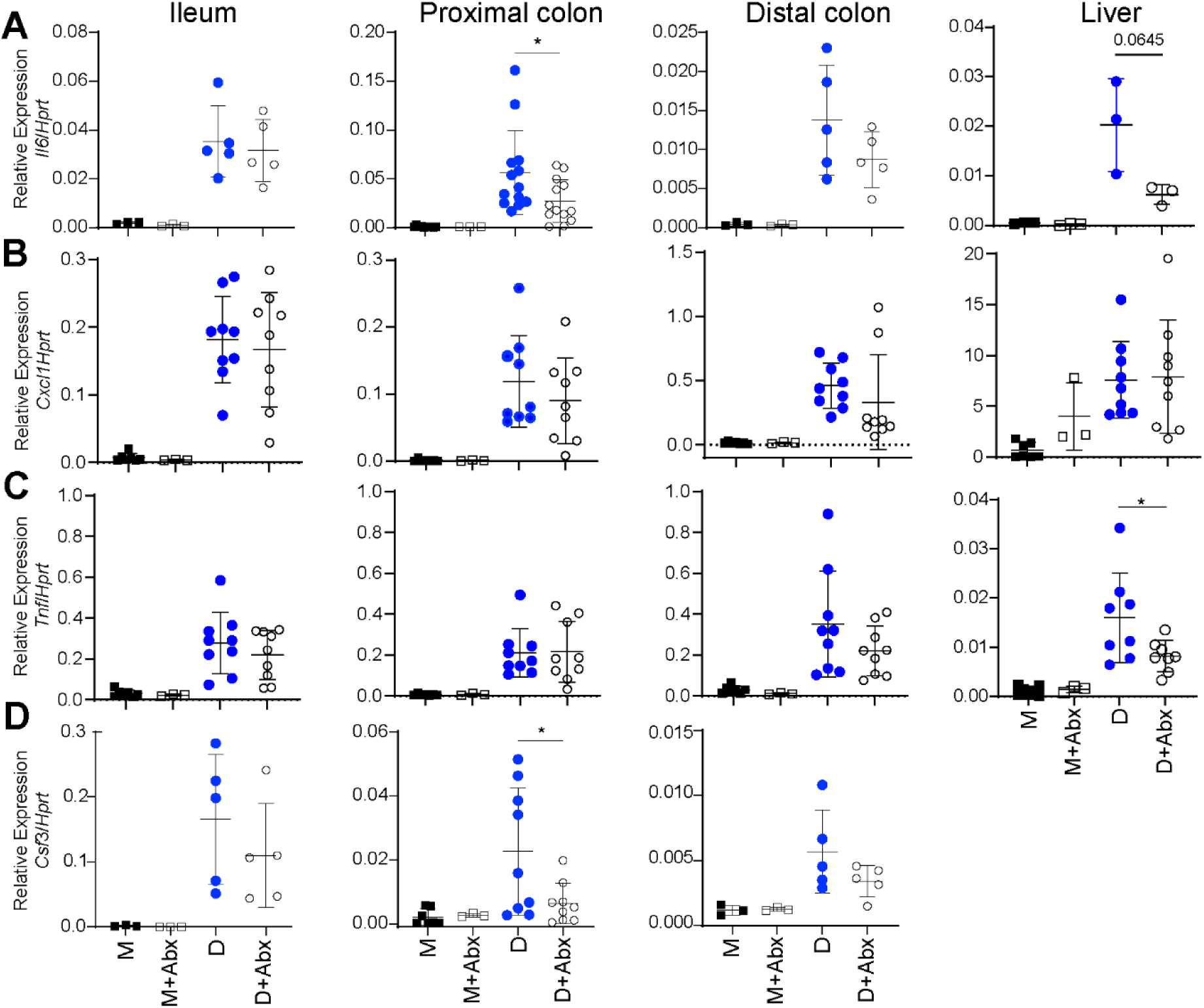
The effect of antibiotics on the expression of mRNAs for cytokines and chemokines in the GI tract and liver during DENV infection. Male AG129 mice were either uninfected or infected with 1×10^5^ FFU DENV2 and treated with or without the 4 antibiotics cocktail from 1 d.p.i as per figure 3. Tissue samples were collected at 4 d.p.i. to assess the levels of mRNAs relative to *Hprt* by qRT-PCR for: A. *Il6*, B. *Cxcl1*, C. *Tnf*, D. *Csf3* (G-CSF), in the indicated tissues, n = 3-13 per group from one to three experiments. Data shows the mean±SD. \**p*<0.05, using an unpaired *t*-test. M, mock-infected; D, DENV-infected; Abx, Abx-treated.

Notably, antibiotic treatment prevented DENV-induced diarrhea (Fig. 3D) and reduced pathology in the colon where bacterial numbers are highest [35, 36] (Fig. 3E; Supp. Fig. 4B). It is worth noting that viremia (Fig. 3F) and circulating NS1 concentrations (Fig. 3G) did not change on day 3 or 4 post-infection although there was a trend to lower viremia on day 4. DENV-induced vascular leak in the liver at 4 d.p.i. showed a non-significant trend for reduction with antibiotics (Fig. 3H). When the antibiotics were used individually, they each had a more modest effect on pathology in the cecum/proximal colon without excessive weight loss (Supp. Fig. 5). Subsequently, a cocktail of three antibiotics - ampicillin, vancomycin and metronidazole – was able to substantially reduce pathology and diarrhea (Supp. Fig 6 and 7).

**Figure 5.**
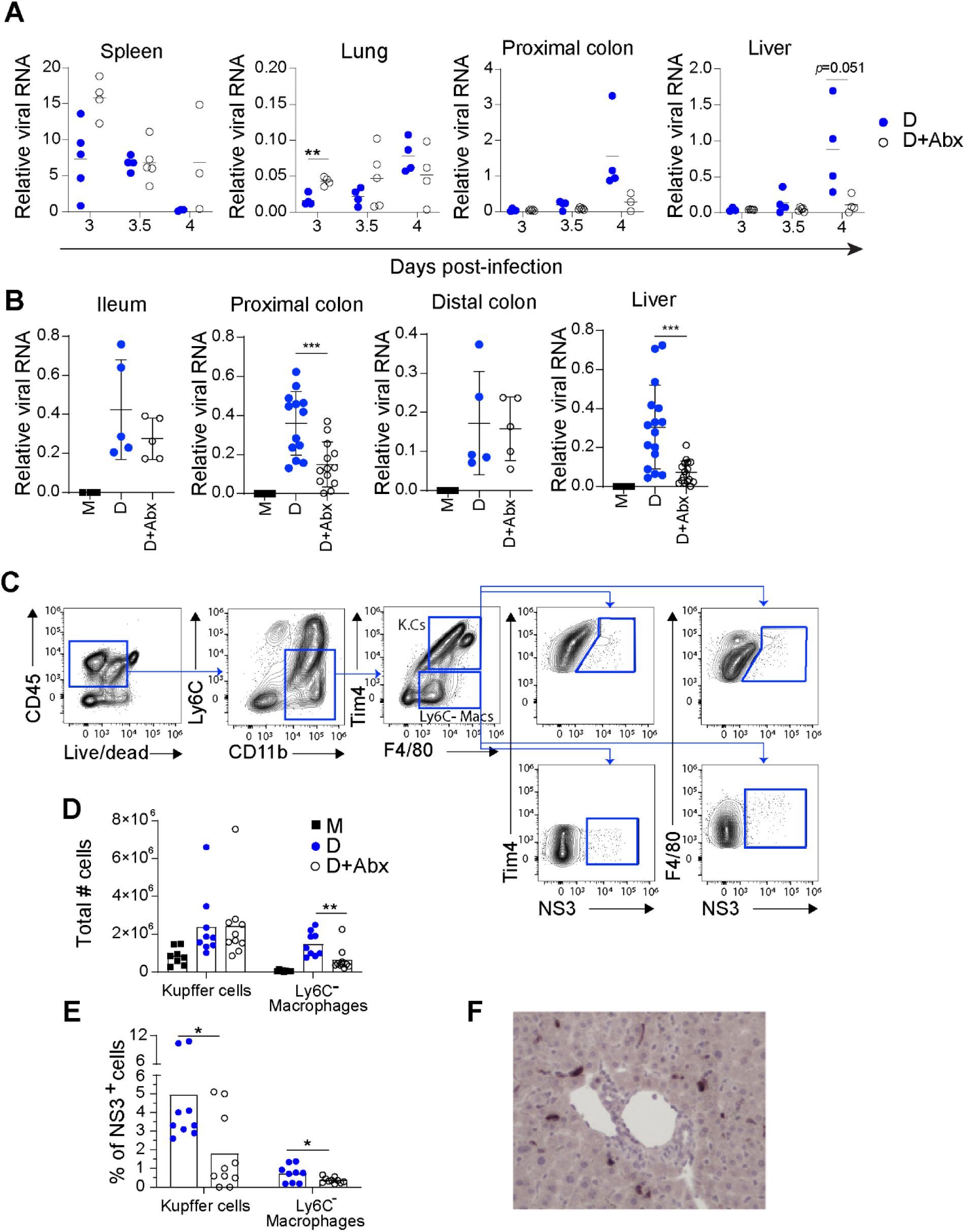
Antibiotic treatment reduces DENV infection in the liver. Male mice were infected with DENV2 as per Figure 3, and treated with 3 (A and D) or 4 (B) antibiotics from 1 d.p.i. **A.** Viral RNA loads relative to mRNA for *Hprt* were assessed in the indicated perfused tissues from 3-4 d.p.i., n = 3-5 per group. **B.** Relative viral RNA levels were assessed in GI-tract and liver at 4 d.p.i., n = 5-16 per group from one to three experiments. Data shows the mean±SD. **C.** Flow cytometry gating strategy for Kupffer cells (CD45^+^ Ly6C^-^ CD11b^+^ F4/80^+^ Tim4^+^) and Ly6C-negative macrophages (CD45^+^ Ly6C^-^ CD11b^+^ F4/80^+^ Tim4^-^). **D.** Total count in whole liver of the indicated cell populations. **E.** Percentage of NS3-positive cells within each cell population. For D and E, n = 8-10 per group from two experiments. \**p*<0.05, \*\**P*<0.01 and \*\*\**p*<0.001 using an unpaired *t*-test. M, mock-infected; D, DENV-infected; Abx, antibiotic-treated. **F.** Immunolabelling of NS3 in liver tissue sections shows infected cells had characteristic Kupffer cell morphology with no hepatocyte infection.

**Figure 6.**
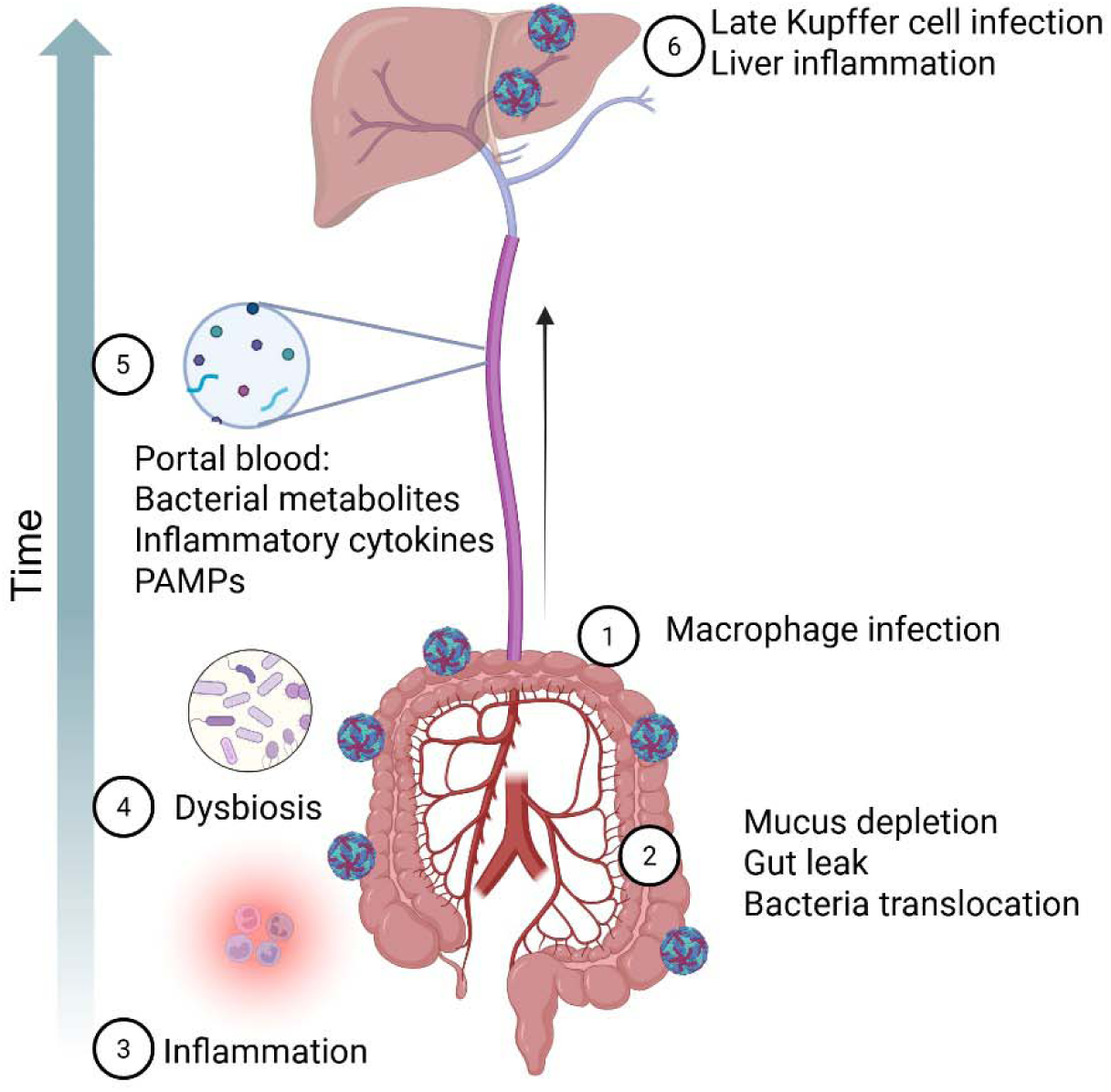
Summary of gastrointestinal features leading to liver damage in severe dengue infection. 1) DENV infected macrophages appear in the small and then large intestine causing **2)** mucus depletion, gut leak, bacterial translocation, **3)** inflammation and **4)** dysbiosis. **5)** Bacteria and gut-derived products including bacterial metabolites, inflammatory cytokines and/or PAMPs are altered by DENV infection and transported directly into the liver via the portal blood. **6)** Some of these products modify the susceptibility of Kupffer cells to DENV infection late in the course of disease and exacerbate liver damage (Figure was created in BioRender https://bioRender.com).

### Antibiotic treatment reduced selected cytokine and chemokine induction in colon and liver

The antibiotic treatment results suggest that gut bacteria and/or bacterial products may play a role in inflammation in the large intestine during DENV infection. We assessed mRNA transcripts of several pro-inflammatory cytokines and chemokines that are pertinent to pathology and homeostasis in the gut (Fig. 4A-D, Supp. Fig. 8). There was a general trend for reduced expression of mRNAs for *Il6*, *Il17f*, *Il22* and *Csf3* (encoding G-CSF) in the colon with antibiotic treatment, and this reached statistical significance in the proximal colon for *Il6* and *Csf3* when sufficient samples were analyzed. Similar to the pathology results (Fig. 3E), there was minimal effect of antibiotics on cytokine levels in the ileum. Expression of mRNAs for *Cxcl1* (Fig. 4B), *Tnf* (Fig. 4C) and *Tgfb1* (encoding TGF-β) (Supp. Fig. 8A) appeared little altered by the antibiotic treatment across the three gut segments. In the liver, expression of mRNAs for *Il6* and *Tnf* were reduced by the antibiotics. Given the effect of antibiotics on gut pathology and cytokine expression, we assessed whether the kinetics of infection and tissue viral load were altered.

### Antibiotics reduce viral RNA load in the proximal colon and liver in the late stages of infection

In prior work we found that viremia followed the kinetics of replication in the spleen, with later appearance of infected cells in the ileum, followed by the colon and liver [11]. The general time course of viral load seen here using perfused tissues was consistent with this, showing high early replication in the spleen and later infection of gastrointestinal tissues such as the proximal colon and liver (Fig. 5A). Unexpectedly, we found that there was a trend for antibiotic treatment to boost levels of virus in the spleen on day 3 as well as the minor amount seen in the lung (Fig. 5A). In contrast, the antibiotics decreased the viral load that appeared at 4 d.p.i. in the proximal colon and liver. This suggested that the effect of antibiotics on reducing viral load might be specific to gastrointestinal tissues and not a direct generalized effect of antibiotics on viral replication (Fig. 5A). Further assessment of gastrointestinal tissues confirmed that viral RNA levels were significantly reduced by the antibiotic treatment in the proximal colon and the liver, with no clear change in the ileum and distal colon at 4 d.p.i., although lower mouse numbers were analyzed (Fig. 5B). The reductions in pathology and inflammatory cytokines in gut tissues seen with antibiotics may thus in part depend on a reduction in viral load but also potentially on reduction in bacteria, which normally increase in abundance along the length of the gastrointestinal (GI) tract [35, 36]. Similar effects on proximal colon and liver viral RNA levels were seen when the three-antibiotic combination was used (Supp. Fig. 7G-H).

The liver receives blood directly from the gut through the portal vein, and the most prominent effect of antibiotic treatment was on reducing the level of viral RNA in the liver, which normally increases on day 4 post-infection (Fig. 5A-B). Kupffer cells, the resident macrophages, are a principal target of DENV infection in the liver of AG129 mice [37] and patients [38]. Using flow cytometry, we confirmed that Kupffer cells (CD45^+^, Ly6C^-^, CD11b^+^, F4/80^+^ and Tim4^+^) and infiltrating Ly6C^-^macrophages (CD45^+^, Ly6C^-^, CD11b^+^, F4/80^+^ and Tim4^-^) are positive for DENV NS3 at 4 d.p.i in AG129 mice (Fig. 5C). The antibiotic treatment did not reduce the total number of Kupffer cells but it reduced the number of infiltrating Ly6C^-^ macrophages into the liver (Fig. 5D). Importantly, the percentage of NS3-positive Kupffer cells and infiltrating Ly6C^-^ macrophages were significantly reduced with the three-antibiotic combination at this late time point of infection (Fig. 5E). We did not observe any infection of hepatocytes or sinusoidal endothelial cells at 4 d.p.i. by flow cytometry or immunohistochemistry (Fig. 5F). We showed here (Fig. 5A) and earlier that the liver was only notably susceptible to infection on day 4 [11] and consequently, it seems that antibiotic treatment is reducing the normal susceptibility of the liver to infection during the late stage of DENV infection. Together with the reduction in liver viral RNA, (Fig. 5A-B) these findings suggest that depletion of the intestinal microbiota limits late-stage Kupffer cell infection and inflammatory cell infiltration into the liver. These data support a model in which gut microbiota-dependent processes enhance liver susceptibility to DENV during severe infection, linking intestinal barrier dysfunction with extraintestinal viral dissemination. This may well be important in the development of severe disease where liver pathology is prominent [39].

## Discussion

Factors promoting severe DENV disease are incompletely understood, but high levels of viremia early in infection and slower viral clearance are implicated [40–42]. The timing has remained puzzling, with serious complications such as hemorrhage, shock and organ damage manifesting with abrupt onset at a time when viremia has declined and fever is abating. However, mouse models show that the decline in viremia and onset of severe disease coincides with the appearance of viral infection in the gut, liver and other organs [11]. The gastrointestinal tract in infected mice shows profound inflammation and pathology [10–12, 43], which may underlie the gut leak observed to correlate with disease severity in patients [5]. Thus, we hypothesize that acute gut inflammatory injury and barrier breakdown leads to an influx of bacterial products, increased systemic inflammation, and disease exacerbation. The results here are generally consistent with this scheme. We detected abrupt onset of gut leak, dysbiosis, and bacterial penetration to the liver. Antibiotic treatment reduced gut pathology and liver viral infection but not viremia, further supporting a role for bacteria in disease amplification rather than primary viral replication. The late rise in viral infection in the liver and the effect of antibiotics in reducing the susceptibility of Kupffer cells to DENV infection is of particular interest due to the importance of liver damage in severe and fatal dengue cases [39, 44–46].

Gut leak has been detected in cases of severe, but not uncomplicated, dengue disease and is associated with elevated circulating microbial LPS and β-glucan [5]. In the mouse model we found variable gut leak, which could be related to a viral load threshold requirement or an effect of variation in the microbiome. Nevertheless, gut leak was confirmed by the presence of live bacteria in the liver and mesenteric lymph nodes. We also have clear evidence of bacterial penetration of the mucosa by silver stain. A prominent histological change in DENV-infected mice is the loss of goblet cell mucus [11], which may be central to barrier dysfunction and exacerbation of intestinal inflammation. Reduced expression of *Muc2* together with diminished expression of goblet cell differentiation factors suggests that impaired goblet cell homeostasis accompanies mucus secretion followed by lack of recovery of cellular mucus content. Similar disruption of the mucus barrier is sufficient to permit bacterial penetration and spontaneous intestinal inflammation in *Muc2^-/-^*mice, supporting a mechanistic role for mucus depletion in DENV-induced gut leak [47–49]. In addition, we document microbiome alterations occurring alongside, or possibly preceding, the detection of DENV-infected cells and inflammatory pathology in the gut [9]. These changes were consistent with dysbiosis, including depletion of putatively beneficial bacteria such as the genera *Limosilactobacillus* and *Lactobacillus*, as well as disruption of normal circadian oscillations in microbial abundance. The early decline in *Lactobacillus* and *Limosilactobacillus* is particularly notable because these taxa promote epithelial barrier integrity, both in humans and mice, through production of antimicrobial metabolites, enhancement of tight junction function and modulation of mucosal immune responses [50, 51]. Whether dysbiosis is a cause or consequence of intestinal inflammation, or both, remains unclear, but our data suggest that disruption of protective microbial communities may contribute to progressive loss of barrier function during DENV infection.

To investigate the role of bacteria in inflammatory pathology and disease severity we treated mice post-infection with a broad-spectrum antibiotic regime. Treatment led to a significant improvement in inflammatory gut pathology, partial restoration of goblet cell mucus content and reduction in diarrhea, along with a reduction in viral RNA in the proximal colon and liver, and fewer DENV-infected Kupffer cells. Effects on inflammation were more apparent in the colon than in the small intestine, correlating with the higher bacterial abundance [35, 36].

Antibiotic treatment caused weight loss over 3 days in uninfected mice, consistent with the established metabolic consequences of microbiome depletion [32–34]. In contrast to our results, a previous study showed that treatment with the same antibiotics for two weeks before *i.v.* infection of *Ifnar1^-/-^*mice with DENV2, West Nile virus (WNV) or Zika virus turned non-lethal infections into uniformly lethal infections [20]. The detrimental effect of pre-treatment with antibiotics may be due to the substantial alteration in hematopoietic progenitors and an impaired T cell response seen in WNV infection [20]. In our experiments, the 3-day treatment post infection may not have been long enough for the effects of changes in haematopoiesis to manifest.

Aggressive treatment with 3-4 antibiotics eliminates most bacteria and is not a realistic therapy. In some countries, up to 17.5% of dengue patients receive antibiotics because of the non-specific disease manifestations and hence difficulties in diagnosis [52]. Antibiotics may also be prescribed to hospitalized dengue patients to prevent secondary bacterial infections [53, 54], although it is reported that antibiotics do not reduce the length of hospitalization [54]. An exception is doxycycline that can directly inhibit DENV replication *in vitro* separate from its antibiotic function [43, 55, 56]. Clinical studies showed doxycycline significantly reduced the length of hospital stay [57], as well as IL-6 and TNF-α levels in serum [58] in severe cases or patients with warning signs. However, given the effect on viral replication *in vitro* the patient improvement may not be due to antibacterial effect.

Recent work implicates a key role for gut bacteria in severe cases of yellow fever virus (YFV) infection. When YFV viremia is clearing, approximately 30% of patients relapse with a return of fever in the intoxication phase. Results from a hamster model, as well as in fatal human cases, suggest that translocation of bacteria from a damaged gastrointestinal tract and subsequent sepsis is responsible for the life-threatening intoxication phase [59]. In both hamsters and human cases, there was no evidence for viral replication in the intestine, and gut damage was suspected to be ischemia-induced necrosis leading to loss of epithelial integrity, profound gut hemorrhage and bacterial influx. This contrasts with the DENV-infection model here, where there is onset of inflammation coincident with the appearance of infected macrophages in the gut, followed by mucus depletion and gut leak without overt disruption of the epithelial layer [11]. Unlike the catastrophic epithelial destruction in YFV infection, our findings indicate a more subtle barrier dysfunction induced by local mucosal DENV infection, leading to an altered hepatic immune microenvironment. Systemic bacterial infections are only confirmed in a minority of severe DENV cases [9].

Vascular leak leading to hypovolemic shock is the key cause of hospitalization with DENV infection, and may be driven by prolonged high viral loads and cytokine storm [13, 40]. Given that circulating LPS is elevated in patients with severe DENV infection [4, 5] and has an established role in vascular leak [4], it is conceivable that gut barrier breakdown is a contributing factor to development of shock. Although dengue shock has sudden onset, vascular leak occurs to a lesser extent even in uncomplicated disease [60], and influx of microbial products may increase cytokine expression above a critical threshold, promoting shock. Although we did not observe a clear decline in vascular leak at 4 d.p.i. with antibiotic treatment, there was a trend towards reduced leak in the liver along with lower *Tnf* and *Il6* mRNA.

In summary, our results demonstrate dysbiosis and gut barrier breakdown in a mouse model of DENV infection, with effects of antibiotics consistent with a role for bacteria in promoting gut inflammatory pathology as well as liver viral load. We propose that the appearance of infected macrophages ([10, 11]) in the intestine initiates inflammation with immune cell infiltrate, cytokine production and goblet cell stress (Fig. 6). Inflammatory damage, dysbiosis, mucus depletion and gut barrier breakdown can generate a self-amplifying loop. These factors are likely to change the composition of the portal blood late in DENV infection, promoting Kupffer cell susceptibility to infection. The consistent effect of antibiotics in decreasing Kupffer cell infection suggests a role for factors such as microbial metabolites or bacteria-dependent gut-derived products in rendering Kupffer cells permissive to DENV. Liver pathology is prominent in post-mortem analysis with infection of both Kupffer cells and hepatocytes noted [45]. *In vitro* infection of human Kupffer cells was reported to be abortive and lead to apoptosis [61]. Kupffer cells act as a firewall protecting the liver from infection, toxins and bacterial products and prominent DENV infection may compromise this function and leave the liver open to hepatocyte infection and damage. Interventions that promote gut barrier integrity and prevent dysbiosis could help to maintain Kupffer cell resistance to infection and reduce liver damage.

## Acknowledgements

The authors thank Jo Gordon for expert histological processing and the UQBR animal house staff; and J. Daniel Bautista for providing the mouse illustration. This work was supported by funding NHMRC project grant 1163313 to K.J.S, H.B-O., and N.M; NHMRC e-Asia JRP grant 2018677 to K.J.S., H.B-O., A.P.Z. and J.K.; NHMRC Ideas grant 2003688 to K.J.S and P.R.V; NHMRC (L1 investigator grant) and Mater Foundation grants to S.Z.H.; and AMED e-Asia JRP grant 25jm0210102h0004 to D.M.T.

