## Supplemental figures for "The gut-liver axis and the role of bacteria in severe dengue virus infection in mice"

**Supplementary Tables**

**Table 1.1 Primers used for quantitative Real-time PCR**

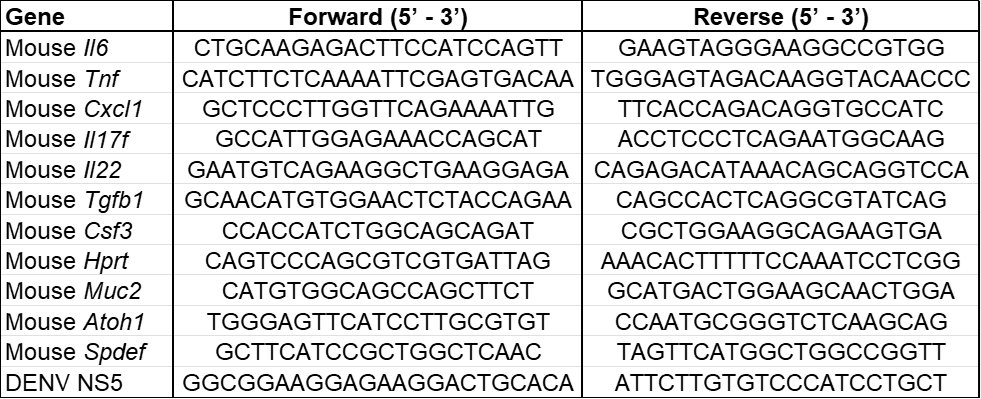

**SUPPLEMENTARY FIGURES**

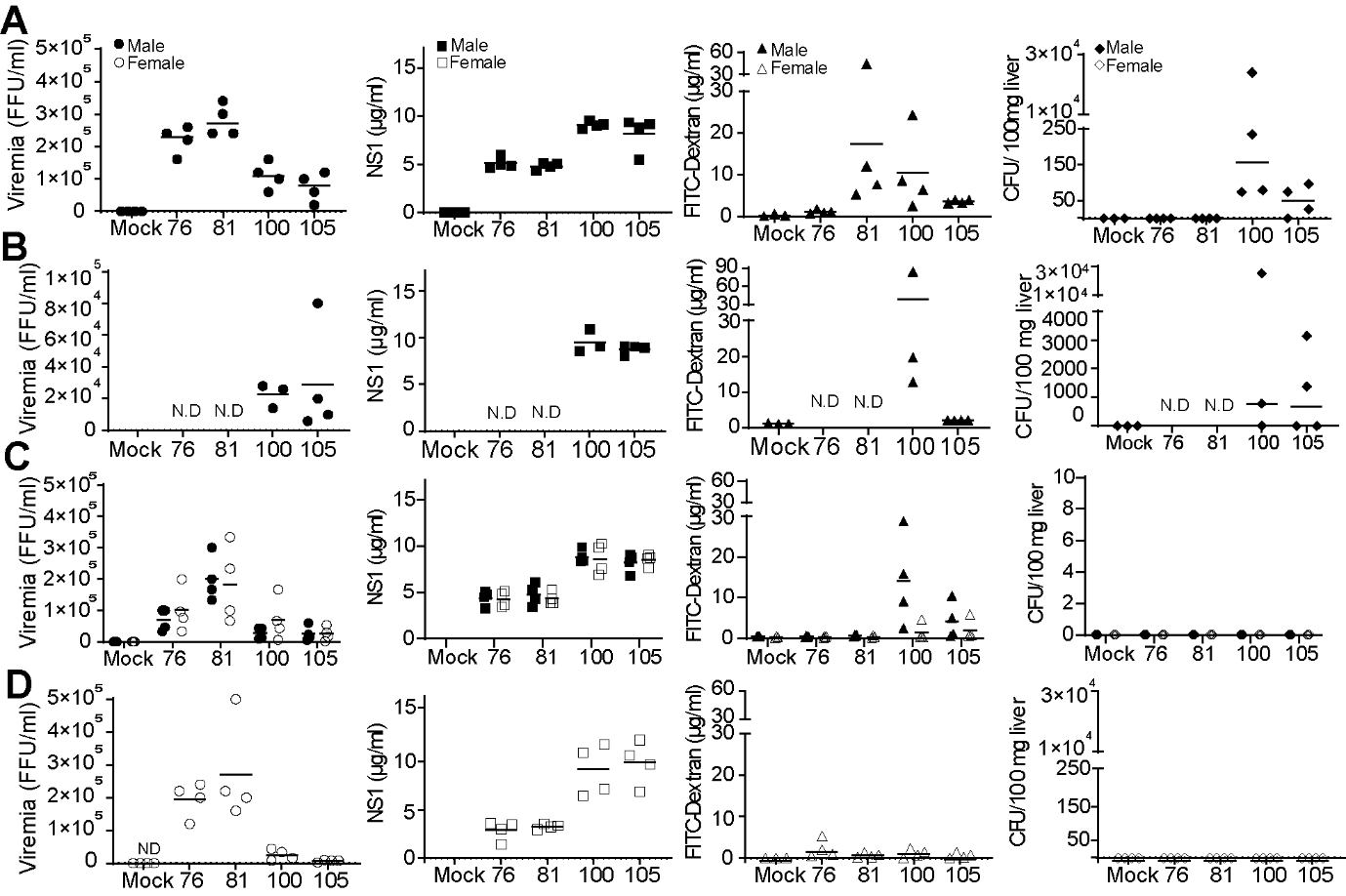

**Supp. Figure 1. Replicate experiments show variable gut leak and bacterial translocation in DENV-infected mice.** Mice were infected with DENV2 and monitored as per Figure 1. A-D show individual experiments with male and female mice indicated by closed and open symbols respectively in all panels. Viremia, levels of NS1, gut leak and bacterial translocation to the liver were assessed at the indicated time points. Gut leak was assessed by detection of 4kDa FITC-Dextran in serum at 4 h post oral intake. Bacteria translocation to the liver was determined by the growth of bacteria under anaerobic conditions from liver homogenates. n = 3 - 4 per group, N.D., not done.

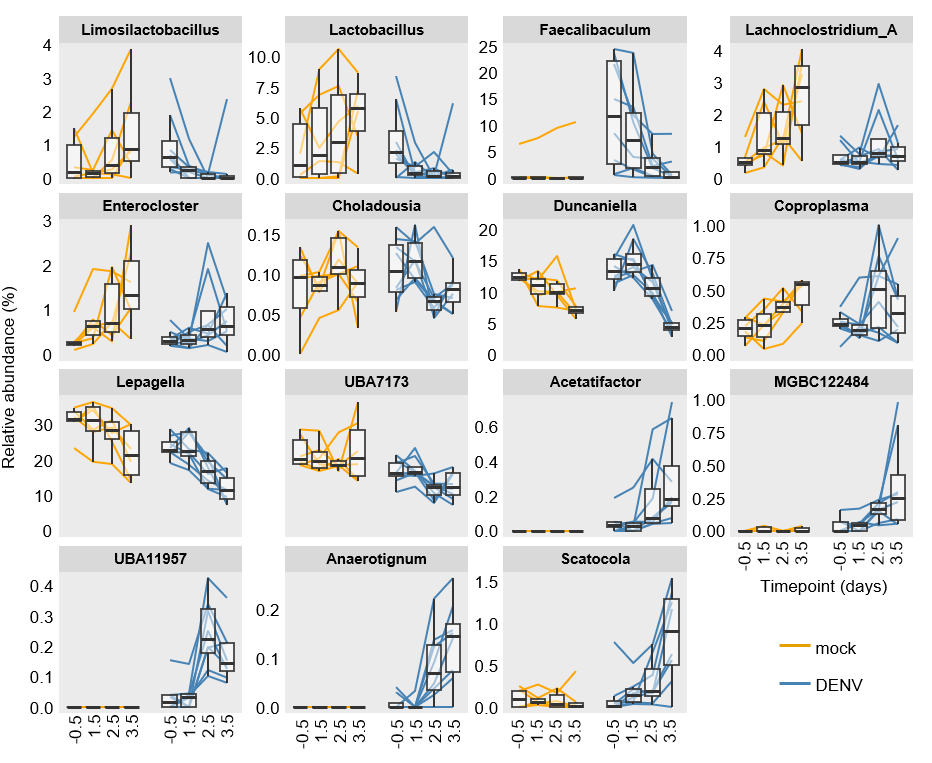

**Supp. Figure 2. Abundances of selected genera**. Relative abundances of genera exhibiting significantly different temporal dynamics between the mock and DENV groups. Boxplots show the distribution of genus abundances, with the central line indicating the median, box edges representing the first and third quartiles, and whiskers extending to the most extreme values within 1.5 × the interquartile range (IQR). Colored solid lines connect longitudinal samples from the same mouse. Each graph corresponds to a different genus, sorted according to the effect size of the treatment-time interaction term shown in the main figure. For some genera, in particular *Faecalibaculum*, differences between mock and DENV-infected groups are evident prior to infection. In addition, changes on day 3.5 occurring in parallel in both groups may be related to stress from blood sampling at day 3.

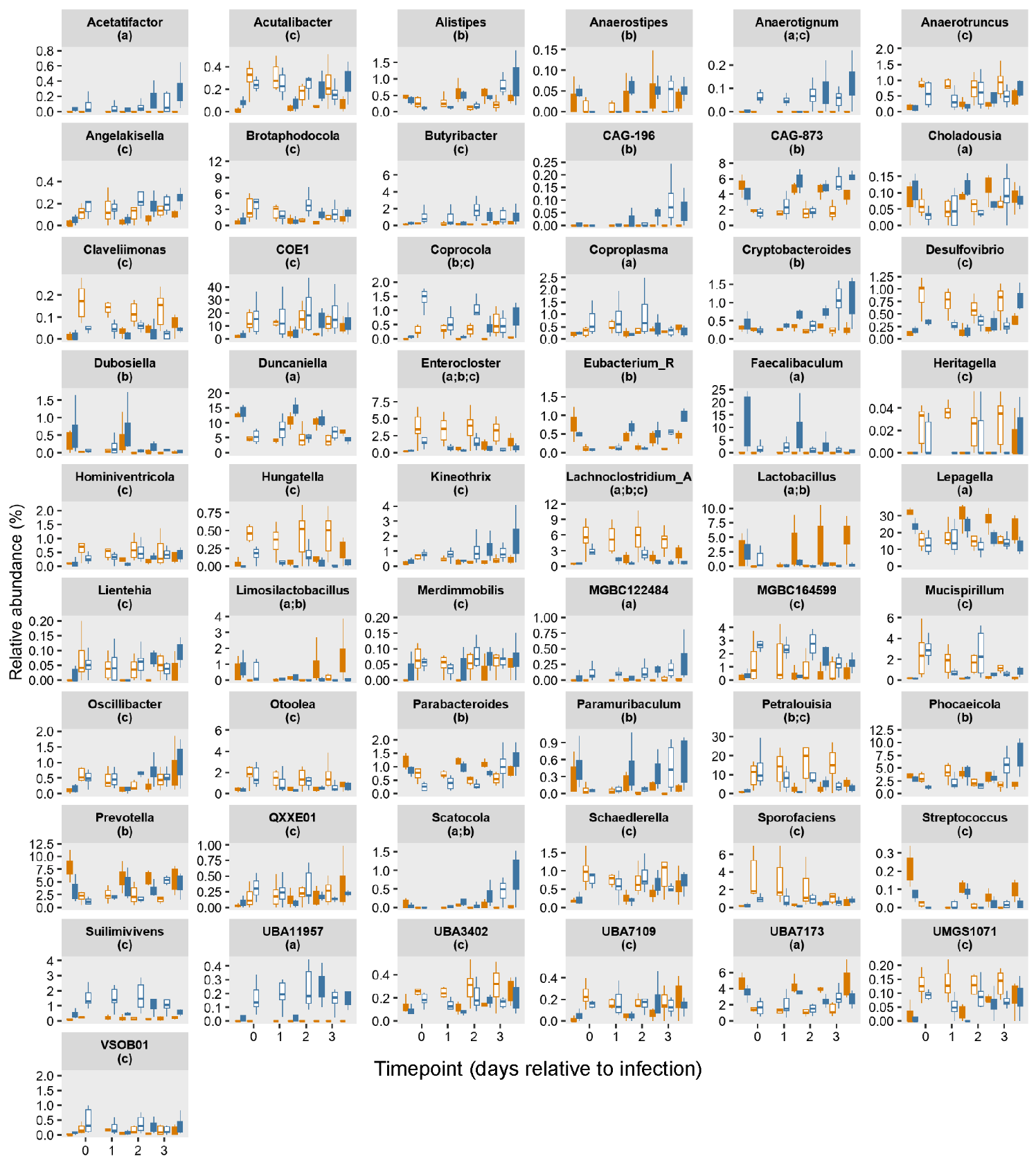

**Supp. Figure 3. Diurnal (circadian) patterns in genus‑level abundances.**

Genus‑level abundances are shown separately for mock‑treated (orange) and DENV‑infected (blue) mice. Boxplots show the distribution of genus abundances, with the central line indicating the median, box edges representing the first and third quartiles, and whiskers extending to the most extreme values within 1.5 × the interquartile range (IQR); outliers are not shown. Colored and white boxes reflect evening and morning samples, respectively. Genera shown represent the union of the following sets: (a) genera with a significant treatment-time interaction term (adjusted P value < 0.05) in evening samples (n = 15); (b) genera with a significant treatment-time interaction term (adjusted P value < 0.05) in morning samples (n = 18); and (c) genera exhibiting highly significant differences between evening and morning samples prior to infection (adjusted P value < 0.001 and |log₂ fold change| > 2; n = 32).

**A**

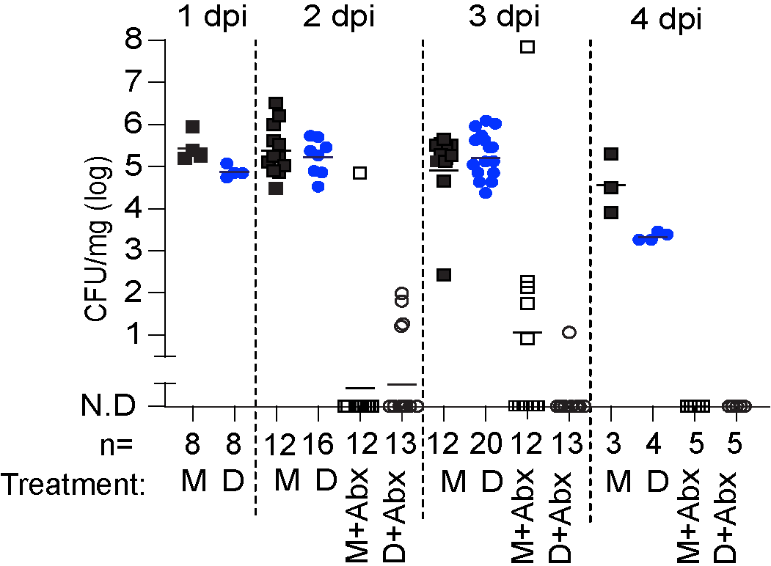

**B**

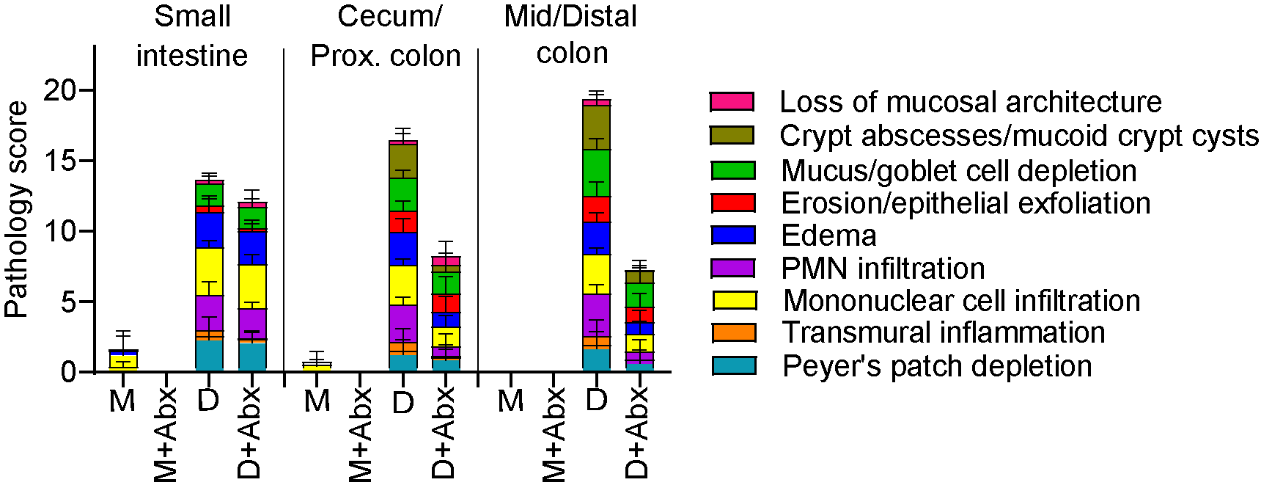

**Supp. Figure 4. Broad spectrum antibiotic cocktail treatment clears fecal bacteria and alleviates substantial gut inflammatory pathology. A.** Reduction of bacteria in the feces cultured under microaerophilic conditions. Each symbol indicates a fecal pellet from one mouse and n for each group is indicated below the axis. There are minimal results at 4 d.p.i. for DENV-infected mice due to diarrhea. **B.** Blinded scoring of gut pathology from two separate experiments shown in Figure 3, n = 11 per DENV-infected group, and n = 10 per mock group. Scores show averaged data from all mice for the 9 parameters assessed. Error bars show SD for each parameter. M, mock-infected, D, DENV-infected, Abx, antibiotic.

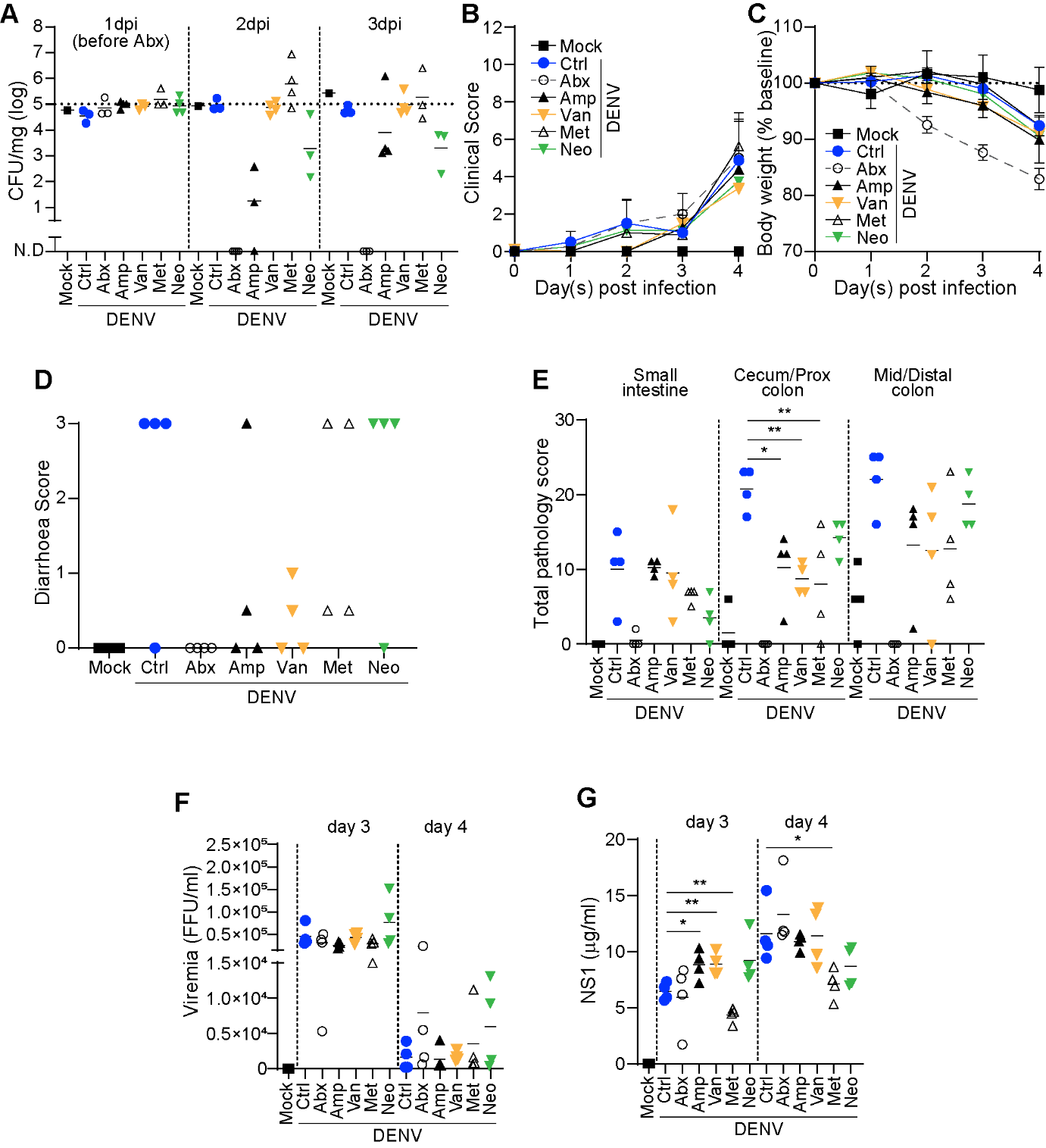

**Supp. Figure 5. Single antibiotics can modestly attenuate gut pathology, with less weight loss than broad spectrum antibiotic treatment.** Eight to 10-week-old male AG129 mice were *i.p.* infected with 1x10^5^ FFU DENV2 and either untreated (Ctrl) or treated with a range of individual antibiotics: Amp, ampicillin; Van, vancomycin; Met, metronidazole; and Neo, neomycin, or a cocktail of all four (Abx) as per Figure 3. **A.** Reduction of bacteria in the faeces grown under aerobic conditions, **B.** clinical scores, **C.** weight loss, **D.** diarrhoea scores at 4 d.p.i., **E.** pathology scores at 4 d.p.i., **F.** viremia, and **G.** levels of NS1 in serum. Lines represent the mean for n = 4 per group from one experiment. B and C are the mean±SD. **p*<0.05, ***p*<0.01 using an unpaired *t*-test. FFU, focus-forming unit.

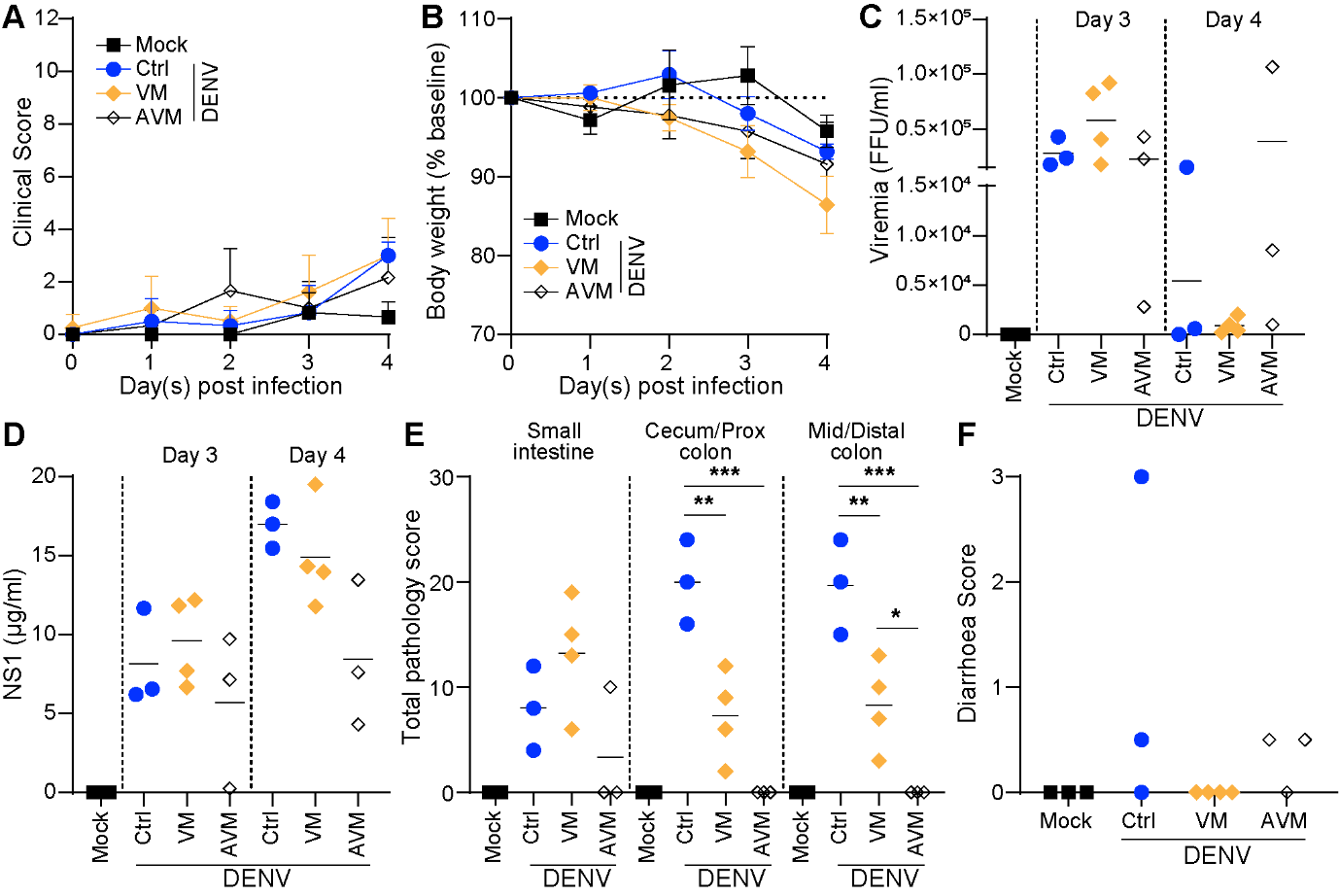

**Supp. Figure 6. A cocktail of ampicillin, vancomycin and metronidazole is more effective than treatment with metronidazole and vancomycin in reducing gut pathology.** Eight to 10-week-old male AG129 mice were *i.p.* infected with 1x10^5^ FFU DENV2. Three or two antibiotics were given in the drinking water as per Figure 3. **A.** Clinical scores, **B.** weight loss, **C.** viremia, **D.** levels of NS1 in serum, **E.** pathology scores at 4 d.p.i. and **F.** diarrhoea incidence at 4 d.p.i. Lines represent the mean and n = 3-4 per group from one experiment. A and B are the mean±SD. **p*<0.05, ***p*<0.01, ****p*<0.001 of unpaired *t*-test. M, mock-infected; D, DENV-infected; FFU, focus-forming unit; VM, vancomycin and metronidazole; AVM, ampicillin, vancomycin and metronidazole.

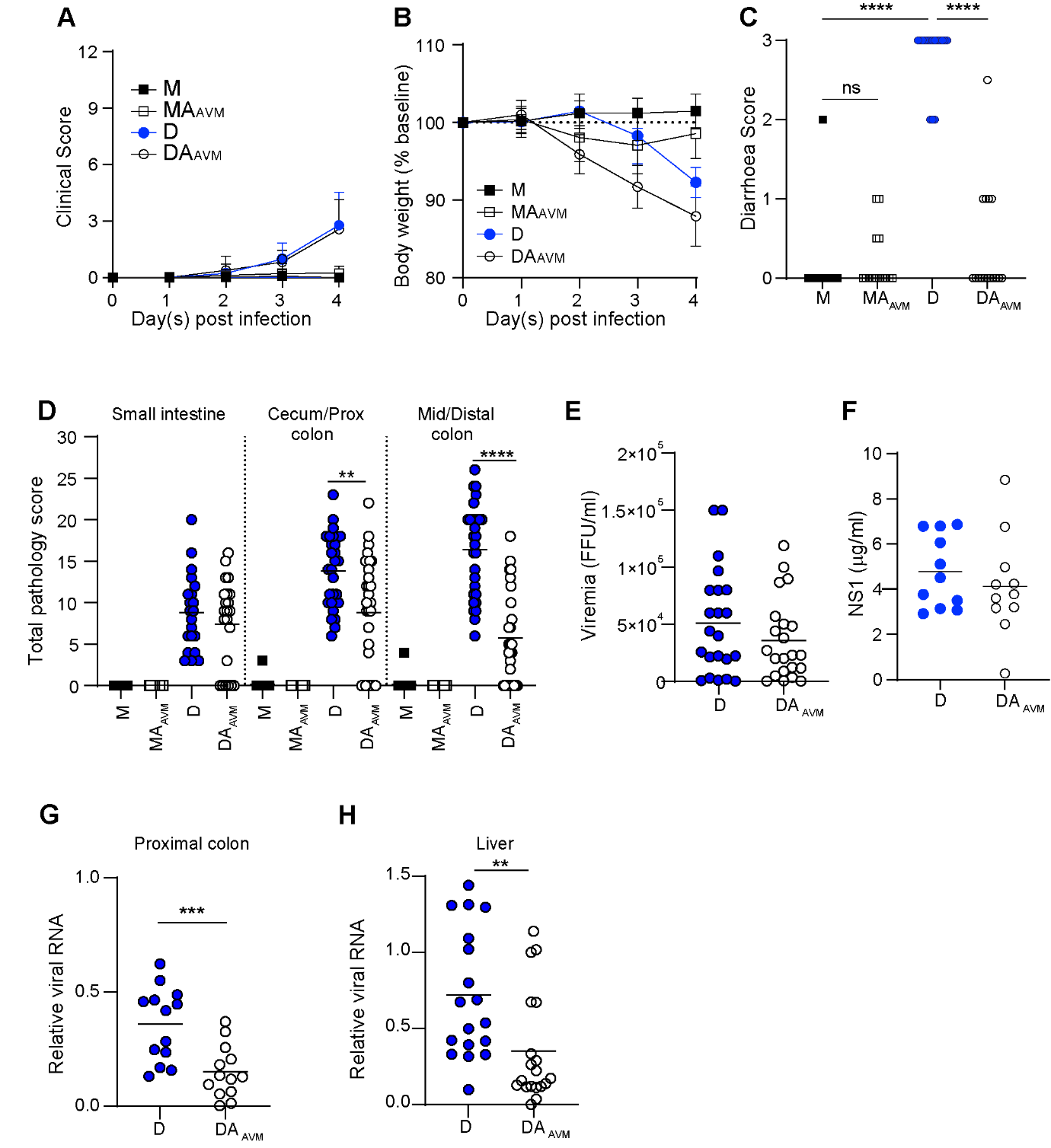

**Supp. Figure 7. A cocktail of 3 antibiotics (AVM) attenuates gut pathology and diarrhoea.** Eight to 10-week-old male and female AG129 mice were *i.p.* infected with 1x10^5^ FFU DENV2. A cocktail of 3 antibiotics (ampicillin, vancomycin and metronidazole) was given in the drinking water from 1 d.p.i. until the end of the experiment. **A.** Clinical scores, **B.** weight loss, **C.** diarrhoea incidence at 4 d.p.i. and **D.** pathology scores at 4 d.p.i. **E** viremia and **F.** levels of NS1 in serum both at 3 d.p.i, and **G. and H.** DENV viral RNA load in the indicated tissues measured relative to *Hprt* mRNA at 4 d.p.i.. **A-C.** n =16-23 per group from five experiments, **D.** n =17-30 per group from six experiments, **E.** n = 22 per group from five experiments, **F.** n = 12 per group from three experiments, and **G. and H.** n = 13-19 per group from four experiments. A and B are the mean±SD, for the others the lines represent the mean, **p*<0.05, ***p*<0.01, ****p*<0.001, *****p*<0.0001 of unpaired *t*-test. M, mock-infected; D, DENV-infected; A_AVM_, antibiotics AVM cocktail-treated; ns, not significant;FFU, focus-forming unit.

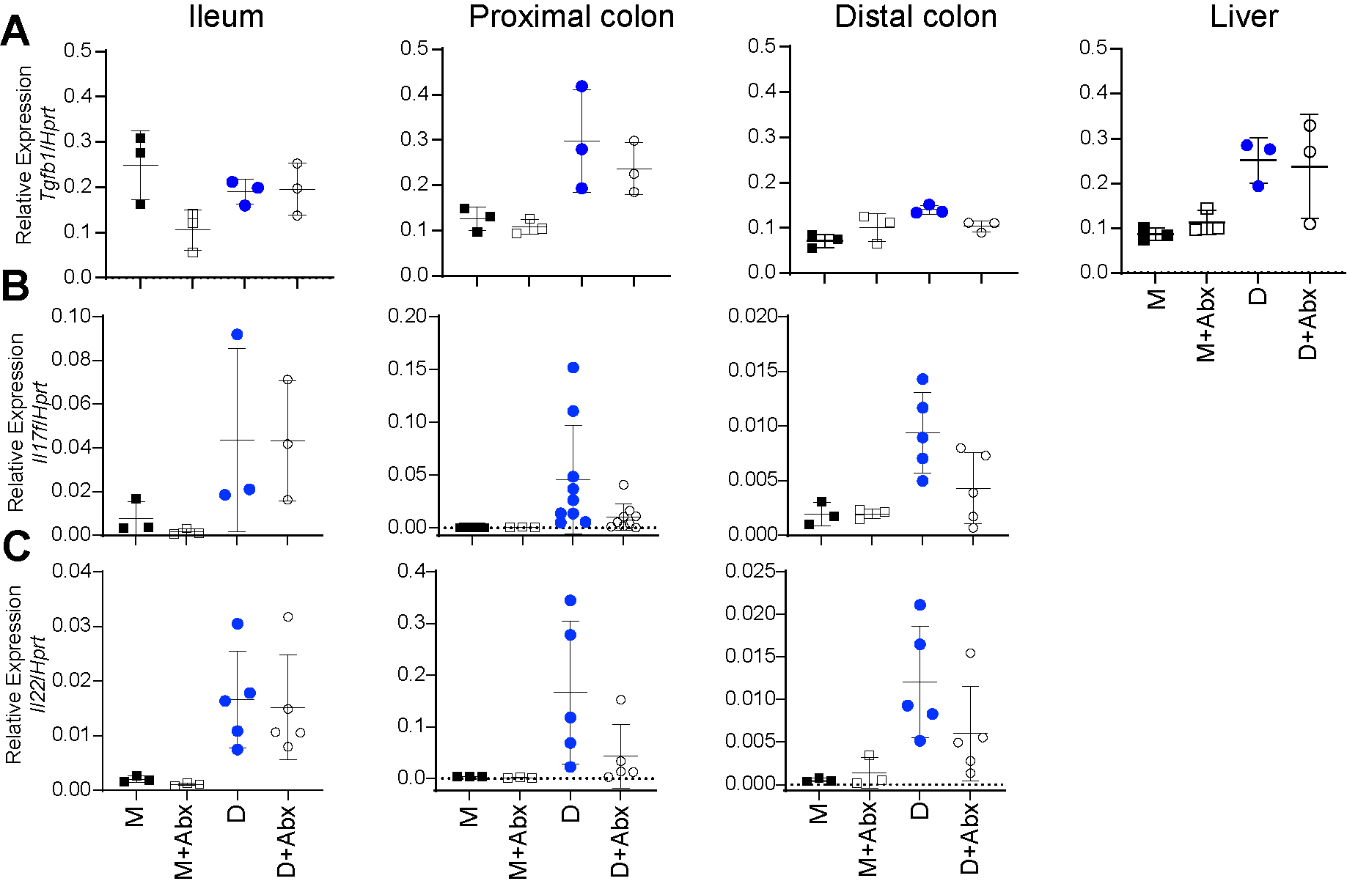

**Supp. Figure 8. The effect of antibiotics on the expression of mRNAs for cytokines and chemokines in the GI tract and liver during DENV infection.** Male AG129 mice were infected with 1x10^5^ FFU DENV2 and treated with or without the 4 antibiotics cocktail from 1 d.p.i as per Figure 3. Tissue samples were collected at 4 d.p.i. to assess the levels of mRNAs relative to *Hprt* by qRT-PCR: **A.** *Tgfb1*, **B.** *Il17f*, **C.** *Il22*, in the indicated tissues. n = 3-9 per group from one to three experiments. Data shows the mean±SD. *p<0.05 of unpaired *t*-test. M, mock-infected; D, DENV-infected; A, Abx-treated.
